# Epiboly in zebrafish requires reactive oxygen species produced by NADPH oxidases for the regulation of vesicular trafficking

**DOI:** 10.1101/2024.12.13.628279

**Authors:** Arlen Ramírez-Corona, Brenda Reza-Medina, Denhi Schnabel, Hilda Lomeli, Enrique Salas-Vidal

## Abstract

Epiboly is the first morphogenetic cell movement that occurs at the onset of gastrulation in zebrafish. During epiboly, the blastoderm thins out and spreads cells over the massive yolk cell. Epiboly progression is controlled by a complex regulatory network that involves diverse molecular effectors. Previously, we reported that reactive oxygen species (ROS) derived from NADPH oxidases (Nox) are required for normal epiboly progression, embryo survival, and early development. We also found that the inhibition of Nox activity during gastrulation downregulates E-cadherin abundance at the enveloping layer (EVL) cell margins. Since the dynamic localization of E-cadherin at the plasma membrane is highly regulated by endocytosis and vesicular trafficking during epiboly, in the present study, we investigated the effects of Nox inhibition and hydrogen peroxide (H_2_O_2_) on endocytosis and in the localization of different proteins important for endosomal trafficking in zebrafish embryos. We show that the simultaneous treatment with the Nox inhibitor VAS2870 and the dynamin 2 (Dnm2) inhibitor dynasore rescues the effects of VAS2870 on epiboly delay, embryo mortality and E-cadherin abundance at EVL cell margins. Furthermore, we found that H_2_O_2_ impacts the endocytic rate of fluorescent fluid-phase markers at the EVL, as well as the localization and abundance of Rab11, a small GTPase protein involved in recycling endosomes. Our results suggest that Nox-derived ROS participate in the regulation of the initial steps of endocytosis and in the endosomal trafficking required for epiboly progression during early zebrafish development.

**HIGHLIGHTS:**

- NADPH oxidase (Nox) activity is required for the epiboly and localization of E-cadherin.
- Dynamin inhibition rescues the developmental defects produced by the loss of Nox activity.
- Nox-derived reactive oxygen species (ROS) participate in the regulation of endosome and E-cadherin trafficking, which is required for epiboly.
- Nox inhibition increases the rate of fluorescent fluid-phase markers of endocytosis in EVL cells.
- H_2_O_2_ decreases fluid-phase internalization in EVL cells.
- H_2_O_2_ regulates Rab11 localization

## INTRODUCTION

Gastrulation is a fundamental morphogenetic process for the early development of metazoans, where massive cell movements participate in the specification of germ cell layers (ectoderm, mesoderm, and endoderm) and in the determination of the embryonic axes (Leptin, 2005). In zebrafish, epiboly is the cell movement that initiates gastrulation. During epiboly, the external epithelial sheet of the blastoderm, known as the enveloping layer (EVL), and the internal deep cell layer (DCL) thin out and spread over the massive yolk cell. In addition, the yolk nuclear syncytium expands vegetally (Warga and Kimmel, 1990). Epiboly onset and progression are regulated by a complex regulatory network (Lepage and Bruce, 2010; Bruce, 2016; Bruce and Heisenberg, 2020). In this context, we described previously that ROS derived from NADPH oxidase (Nox) catalytic activity are required for blastoderm cell motility, promoting epiboly progression and embryo survival (Mendieta-Serrano et al., 2019). To inhibit Nox activity, we used VAS2870, a cell-permeable triazolo pyrimidine with no reported antioxidant activity (ten Freyhaus et al., 2006; Altenhofer et al., 2012). Treatment of zebrafish embryos with VAS2870 affects key components that control and promote epiboly progression, such as the actin and microtubule cytoskeleton. We also observed a significant decrease in E-cadherin (E-cad) at EVL cell boundaries after the inhibition of Nox activity, an effect that was reversed by H_2_O_2_ treatment (Mendieta-Serrano et al., 2019). E-cad belongs to the protein superfamily of cadherins, which are membrane proteins that mediate homotypic intercellular adhesion in animal cells (Nose et al., 1990; Shapiro et al., 1995) and are required for epiboly during normal development (Kane et al., 1996; Babb and Marrs, 2004; Kane et al., 2005). Early reports revealed that disruption of E-cad-dependent adherens junctions by calcium chelating treatment results in E-cad endocytic uptake along with fluid-phase markers (Kartenbeck et al., 1991) and can be recycled back to the plasma membrane, similar to other membrane proteins (i.e., transferrin receptor) (Le et al., 1999). The internalization and trafficking of E-cad are regulated by clathrin, dynamin 2, Rab5 and Rab11 (Le et al., 1999; Lock and Stow, 2005), among many other proteins.

Different signaling cascades modulate endosomal trafficking and regulate E-cad persistence at the plasma membrane (Song et al., 2013; Sempou et al., 2016). However, as mentioned above, we showed that the decrease in ROS decreases E-cad abundance at EVL cell margins (Mendieta-Serrano et al., 2019). These results suggest that ROS might be important for the regulation of endocytosis and endosome trafficking, and that the observed changes in the abundance of E-cad in the EVL plasma membrane in VAS2870-treated embryos can be used as a readout in the analysis of the participation of Nox-derived ROS in the regulation of endocytosis.

In the present work, we found that the delay in epiboly progression and the decrease in embryo survival caused by VAS2870 are reversed by the pharmacological inhibition of endocytosis by dynasore treatment. We also found that VAS2870 increases the uptake of the fluorescent fluid-phase marker dextran-Alexa, while treatment with hydrogen peroxide (H_2_O_2_) decreases it. The pharmacological inhibition of endocytosis by dynasore treatment also rescued the membrane E-cad signal at the EVL cell margins in VAS2870-treated embryos, indicating that endocytosis inhibition by dynasore increased the retention of E- cad at the plasma membrane. Finally, we also found that VAS2870 alters the localization of Rab11, a small GTPase involved in the process of vesicle recycling, an effect that is rescued by dynasore or H_2_O_2_ treatment, stressing the relevance of ROS in the regulation of endocytosis and vesicular trafficking during early development.

## MATERIALS AND METHODS

### Fish maintenance and strains

Wild type zebrafish (*Danio rerio*) AB embryos obtained from the Zebrafish International Resource Center (ZIRC) and embryos from a strain established in the laboratory were raised by standard procedures and developmental stages were selected by morphological criteria (Kimmel et al., 1995). Zebrafish were handled in compliance with local animal welfare regulations and approved by the Institute’s Ethical Committee (Instituto de Biotecnología, UNAM).

### Chemical treatment of zebrafish embryos

VAS2870 was obtained from different distributors (SML0273, Sigma-Aldrich; 19205, Cayman Chemical; 6654, Tocris; sc-471103 Santa Cruz), hydrogen peroxide (H1009, Sigma-Aldrich), dynasore (D7693, Sigma-Aldrich), MG-132 (C2211, Sigma-Aldrich) chloroquine (C6628, Sigma-Aldrich), and ammonium chloride (A9434, Sigma-Aldrich). Absolute anhydrous DMSO (276855, Sigma-Aldrich) was used to generate stock solutions of the inhibitors. Fertilized eggs were incubated at 28 °C in embryo rearing medium (ERM) (Westerfield, 2000) until reaching the desired developmental stage to start treatments. Treatments used were 10 µM VAS2870, 1 mM and 2.5 mM H_2_O_2_, 75 µM dynasore, 50 µM chloroquine, 10 mM ammonium chloride (NH4Cl) and 5 µM MG-132, either individually or combined in rescue assays. Control embryos were treated with 1% DMSO (v/v) in ERM and processed as described below. Embryos in their chorions were used in all the experimental procedures, unless otherwise indicated. Experiments were replicated at least three times. The total number of embryos per experimental treatment are indicated in the figure legends.

### Immunofluorescence for E-cadherin, ZO-1, clathrin, dynamin 2, Rab11, F-actin and nuclei detection

Whole-mount staining for F-actin cytoskeleton, nuclei and immunofluorescences in zebrafish embryos were performed as previously described (Mendieta-Serrano et al., 2019). The primary antibodies used were mouse anti-E-cadherin at 1:250 (610182 [AB_397581], BD Biosciences) rabbit anti-Rab11a/b at 1:200 (GTX127328, [AB_11171170] Genetex), rabbit anti-Clathrin at 1:200 (ab59710, [AB_941047] Abcam), rabbit anti-Dynamin 2 at 1:100 (GTX127330, [AB_11163665] Genetex) (Eno et al., 2016) and mouse anti-ZO-1 at 1:200 (33-9100, [AB_2533147] Invitrogen) (Sonal et al., 2014). The secondary antibodies employed were goat anti-mouse Alexa 647 at 1:100 (A21235 [AB_2535804], Invitrogen) and goat anti-rabbit Alexa 647 at 1:100 (A21244, [AB_2535812], Invitrogen). Actin filaments were stained with Alexa fluor 488 phalloidin at 1:100 (A12379, Invitrogen) and nuclei with DAPI at 1:3000 (D3571, Invitrogen).

### Confocal microscopy images acquisition

Embryos were imaged with the Marianas 3i spinning disk confocal microscope with a CSU-W1 unit (Yokogawa Electric Corporation) mounted on an inverted Observer Z.1 microscope (Zeiss), with an iXon EMCCD camera (Andor Technology). Epiboly progression was evaluated from images taken with a 10X objective (numeric aperture 0.3), and immunofluorescence images were taken with a 63X objective (numeric aperture 1.4). Depending on the fluorescent dyes used, the embryos were excited with a wavelength of 405 nm, 488 nm or 640 nm. Confocal parameters like camera intensification and exposure times were maintained constant during image acquisition for quantitative comparisons among treatment groups. All image stacks were exported as Tiff format image-series with the 3i SlideBook Software (www.intelligent-imaging.com) for further analysis using Fiji software (Schindelin et al., 2012).

### Epiboly progression and fluorescence microscopy analysis

For a detailed analysis of epiboly progression of the EVL, DCL and YCL nuclei, fixed and dechorionated embryos were double stained with DAPI and Alexa fluor 488 phalloidin and imaged in lateral view as previously described (Mendieta-Serrano et al., 2019) with slight modifications. Maximum intensity projection images of the different channels were obtained with the Fiji software (Schindelin et al., 2012). The percentage of epiboly progression of the different cell layers was measured by comparing the distance between the animal pole and the edge of migration of each layer against the total diameter of the embryo, taken from the animal to vegetal pole.

### Survival assays

Treatments with the different inhibitors started at sphere stage and embryos were incubated 24 hrs at 28 °C. The next day the surviving embryos were counted, dechorionated and transferred into 35 mm dishes covered with 1% agarose prepared with ERM for imaging. Embryos were anesthetized using 168 µg/ml (w/v) of Tricaine (ethyl 3-aminobenzoate methanesulfonate) (E10521, Sigma) in ERM. Images were taken with an Axiocam 208 color camera from Zeiss. For survival curves using chloroquine, NH4Cl and MG-132, the number of surviving embryos was counted at sphere stage, shield stage, 70% and 90% epiboly, tailbud and 24 hpf.

### Enveloping layer E-cadherin, clathrin, dynamin 2, Rab11, ZO-1 and F-actin quantification

EVL cell margin masks were done as previously described (Mendieta-Serrano et al., 2019), using the F-actin maximum intensity projections images from confocal images of EVL cells and the Fiji software (Schindelin et al., 2012) . Images were processed with a medium blur filter with a radius of 2 pixels, and EVL cells were outlined using the Morphological segmentation plugin (Legland et al., 2016). A distance map of the resulting watershed images and thresholding with a value of 5 were used to obtain the F-actin binary images with outlined EVL cells. These binary images were used to generate the cell area and the cell margin masks.

The E-cad signal located at the cell margins or in cytoplasmic vesicular structures was also quantified using Fiji software. To quantify the number of vesicle structures positive for E- cad, we used a threshold segmentation with the intermodes method in single confocal images. The cell masks were used to select only the segmented elements inside of the EVL cell areas and not membranes. The number of vesicles was calculated with the Analyze Particles command. To quantify the E-cad, clathrin, dynamin 2 and ZO-1 levels at EVL cell margins, we used margin cell masks to measure the mean fluorescent intensity in maximum projection images. The mean fluorescent intensity of clathrin, dynamin 2 and Rab11 was also measured inside the cytoplasm of EVL cells.

Rab11 fluorescent signal at the EVL cell margins was quantified by using maximum projection images and a threshold segmentation with the intermodes method to generate binary images of the fluorescent signal. The number of segmented fluorescent pixels at the cell margins was counted with the histogram tool (pixels with a grey value of 255).

### Enveloping layer microridge protrusions quantification

Microridge protrusions are actin rich structures at the EVL cells (Inaba et al., 2020) and they were quantified as previously described (Mendieta-Serrano et al., 2019). Confocal images of control and treated embryos stained with Alexa fluor 488 phalloidin were processed with the Unsharp Mask filter. Afterwards, EVL cell margins were segmented using the Morphological segmentation plug in (Legland et al., 2016) and actin structures were segmented with the intermodes method and quantified with the Analyze particles plugin (Bolte and Cordelieres, 2006). Microridges were quantified either per cell or per area to take into consideration changes in cell area due to the treatment with VAS2870.

### Fluid-phase endocytosis assay

This procedure is a modified version from protocols described by others (Lepage et al., 2014; Willoughby et al., 2021). In brief, embryos were manually dechorionated at sphere stage and placed in a 48 well plates with 10 embryos per well. Treatments were administered with 300 µl of media per well. Wells were previously coated with 1% agarose in ERM medium to prevent embryos from sticking to the bottom. One hour after the beginning of treatment with the drugs, dextran 10,000 MW coupled with Alexa fluor 594 (D22913, Invitrogen) was added to the media at a concentration of 0.5 µg/ml (w/v). Embryos were incubated with the drugs and fluorescent dye for 2 hrs at 28 °C covered form the light, afterwards washed with ERM, fixed with 4% PFA in PBS and then processed as previously described to stain nuclei, actin cytoskeleton (Mendieta-Serrano et al., 2019) and ZO-1 (Sonal et al., 2014). Embryos were excited with a wavelength of 405 nm, 488 nm, 561 nm or 647 nm depending of the fluorophore, and imaged as described above. Quantification of dextran florescent signal was carried out by stacking images containing EVL cells and creating maximum intensity projections. Fluorescent signal was also segmented by threshold with the triangle method and vesicles were counted with the Analyze particles plugin.

## Statistical analysis

No statistical method was used to predetermine sample size. In each experiment, embryos were randomly selected, and the investigators were not blinded to allocation during experiments and outcome assessment. No outlier test was performed to exclude embryo samples from the analysis. Statistical analysis was performed with three independent experimental replicates, except where indicated. Statistical analysis was performed with Prism (GraphPad). Normality tests were carried out depending on the number of analyzed samples: D’Agostino, Shapiro-Wilk or Kolmoronov-Smirnov. Data was compared using analysis of variance (Ordinary ANOVA or Kruskal-Wallis) with Tukey’s or Dunn post-hoc analysis. Error bars in boxplots represent the minimum and maximum values within the population. Mann Whitney or Welch’s t test were also used to compare treatments.

For survival analysis we performed a Gehan-Breslow-Wilcoxon test to compare the survival curves between the different treatment groups.

## RESULTS

### Dynamin inhibition rescues delayed epiboly and embryo lethality caused by Nox activity downregulation

In our previous report, we showed that Nox-derived ROS are required for blastoderm cell motility during epiboly in zebrafish embryos (Mendieta-Serrano et al., 2019). To decrease ROS formation, we used the general Nox pharmacological inhibitor VAS2870 (ten Freyhaus et al., 2006). Embryos in their chorions were incubated in control media, ERM (Westerfield, 2000) with 1% (v/v) dimethyl sulfoxide (DMSO), to permeabilize the chorion (Kais et al., 2013). The same media was used as a vehicle for treatments with different inhibitors. To decrease Nox activity, the embryos were treated continuously with 10 μM VAS2870, from the oblong-sphere stage and throughout epiboly progression. As previously shown, in these embryos, EVL and DCL epiboly progression is significantly delayed compared with that in control-treated embryos (Figure 1A to A”, B to B” and D to D’). In contrast, YSL nuclei showed no significant epiboly delay compared with those of control embryos (Figure 1B’ and D”). These effects on epiboly caused by VAS2870 exposure are somewhat similar to the reported effects on epiboly observed when different adhesion molecules are knocked down by morpholinos. For example, prion protein 1 (Prp- 1) morphant embryos show a significant epiboly delay, and E-cad turnover is also severely affected. Interestingly, in these morphants, the epiboly delay and E-cad defects can be rescued by exposure to dynasore (Sempou et al., 2016), a dynamin-dependent endocytosis inhibitor that interferes with vesicle fission (Macia et al., 2006). Additionally, the inhibition of proteasomal degradation with MG-132 (Palombella et al., 1994) significantly rescued epiboly and E-cad localization in Prp-1 morphants, whereas lysosome inhibition with chloroquine (Homewood et al., 1972; Redmann et al., 2017) or with ammonium chloride (NH4Cl) (Hart et al., 1983) produced only a partial improvement (Sempou et al., 2016).

**Figure 1.**
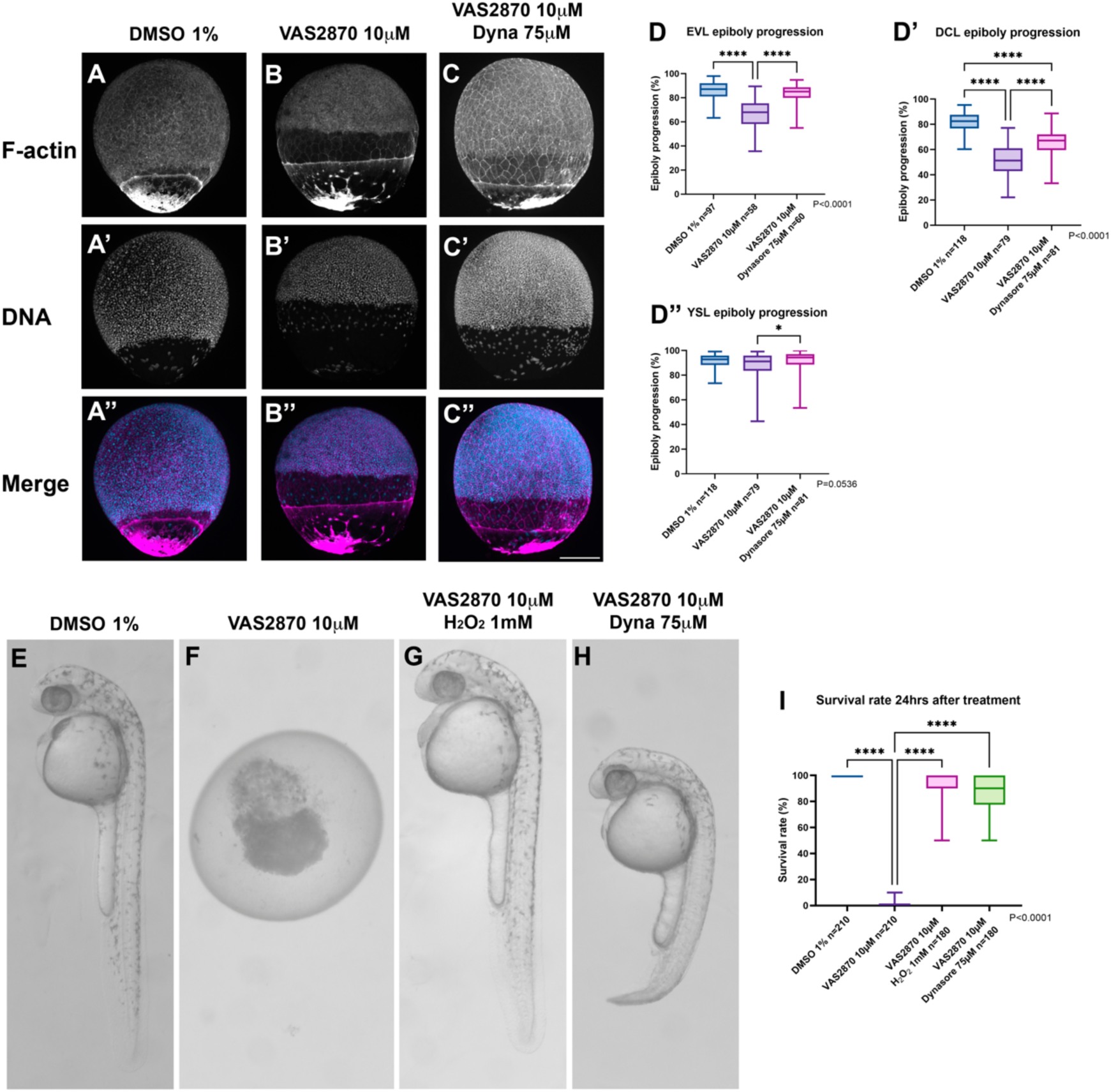
Inhibition of endocytosis restores epiboly, embryo survival and morphology affected by Nox activity down regulation. (A-C’’) All treatment groups started with embryos at oblong-sphere stage and the experiment stopped when control treatment embryos reached around 80% epiboly. Embryos were then fixed, stained with DAPI and phalloidin-Alexa and visualized in a spinning disk confocal microscope. (A-A”) Control embryos were incubated with vehicle (1% DMSO). (B-B”) Embryos treated with 10 µM VAS2870 show a delayed epiboly progression. (C-C”). Delayed epiboly progression due to VAS2870 exposure is rescued by treatment with 75 µM Dynasore (Dyna). Scale bar, 200 µm. (D-D”) Enveloping layer (EVL), Deep cell Layer (DCL) and Yolk Syncytial Layer (YSL) epiboly progression quantification in the different treatment groups. Values represent median and interquartile range; whiskers show minimum and maximum values.\**p* < 0.01 by Kruskal-Wallis one-way analysis of variance (ANOVA) with Dunns’s post hoc analysis (*n*, indicated in the graph). N=3 experimental repetitions. (E-H) All treatment groups started with embryos at oblong-sphere stage, embryo survival and morphology were analyzed 24hrs after pharmacological treatments (embryos around prim-16/31 hpf). (E) Embryos treated in control media (1% DMSO). (F) Embryos treated with 10 µM VAS2870. (G) Embryos treated with 1 mM H_2_O_2_ and VAS2870 simultaneously. (H) Embryos treated with VAS2870 and 75 mM Dynasore simultaneously. Scale bar, 300 Scale bar, 300 µm. (I) Quantification of embryo survival 24hrs post treatment. Values represent median and interquartile range; whiskers show minimum and maximum values. \**p* < 0.01 Kruskal-Wallis one-way analysis of variance (ANOVA) with Dunns’s post hoc analysis (*n*, indicated in the graph). N=3 experimental repetitions.

On the basis of these findings, we analyzed whether the downregulation of endocytosis by dynasore treatment is capable of rescuing VAS2870-induced defects.

First, we found that the exposure of embryos to 75 μM dynasore, did not affect epiboly (Supplementary figure 1B to B” and E to E”). Remarkably, the epiboly delay of both EVL and DCL elicited by VAS2870 treatment was significantly rescued by simultaneously treating embryos with 75 μM dynasore (Figure 1C to C”, and D to D”). These results suggest that VAS2870 and dynasore pharmacological treatments compensate for each other’s effects on EVL and DCL epiboly.

We also reported that zebrafish embryos continuously exposed to VAS2870 die before 24 hours post fertilization (hpf), and the addition of exogenous H_2_O_2_ rescues embryo survival and development (Mendieta-Serrano et al., 2019) (Figure 1 E to G, and I). Since endocytosis inhibition with dynasore rescued the epiboly delay caused by VAS2870 exposure, we investigated whether this treatment also rescued embryo survival and development. Simultaneous treatment with VAS2870 and dynasore not only rescued embryo survival by 24 hpf (Figure 1H and I) and 48 hpf (Supplementary Figure) but also allowed development to proceed. Approximately 50% of the embryos develop normally, approximately 50 % show developmental delay, and 13 % of those in the delayed group display normal caudal morphology (Supplementary Figure 4). These findings indicate that the inhibition of dynamin-dependent endocytosis is sufficient to successfully rescue the development of zebrafish embryos lacking Nox activity.

### Lysosome or proteasome inhibition fails to rescue embryo survival after ROS downregulation with VAS2870 treatment

Given that the delaying effects of Nox inhibition on epiboly can be rescued by pharmacologically inhibiting dynamin-dependent endocytosis, it is possible that in VAS2870-treated embryos, the increase in endocytosis led to increased protein degradation by the lysosome or proteasome system.

To assess this possibility, we tested whether the inhibition of any of these protein degradation systems compensated for the effects of Nox-derived ROS downregulation on epiboly and embryo survival. First, we tested different concentrations of the lysosome inhibitors, chloroquine and NH4Cl, and the proteasome inhibitor MG-132, to determine the highest concentration at which each of these inhibitors can be used without affecting early development (not shown). We subsequently tested the effects of each inhibitor on embryos exposed to VAS2870.

We found that simultaneous treatment with VAS2870 and 5 μM MG-132 did not significantly affect epiboly progression after Nox inhibition (Figure 2C to C” and E to e”). Similarly, lysosome inhibition with 50 μM chloroquine (Figure 2D to E” and F to F”) or 10 mM H4ClN (Supplementary Figure 2) failed to rescue epiboly progression in both blastoderm cell layers. These results indicate that Nox-derived ROS participate in the regulation of protein localization dynamics but apparently not in the regulation of protein degradation.

**Figure 2.**
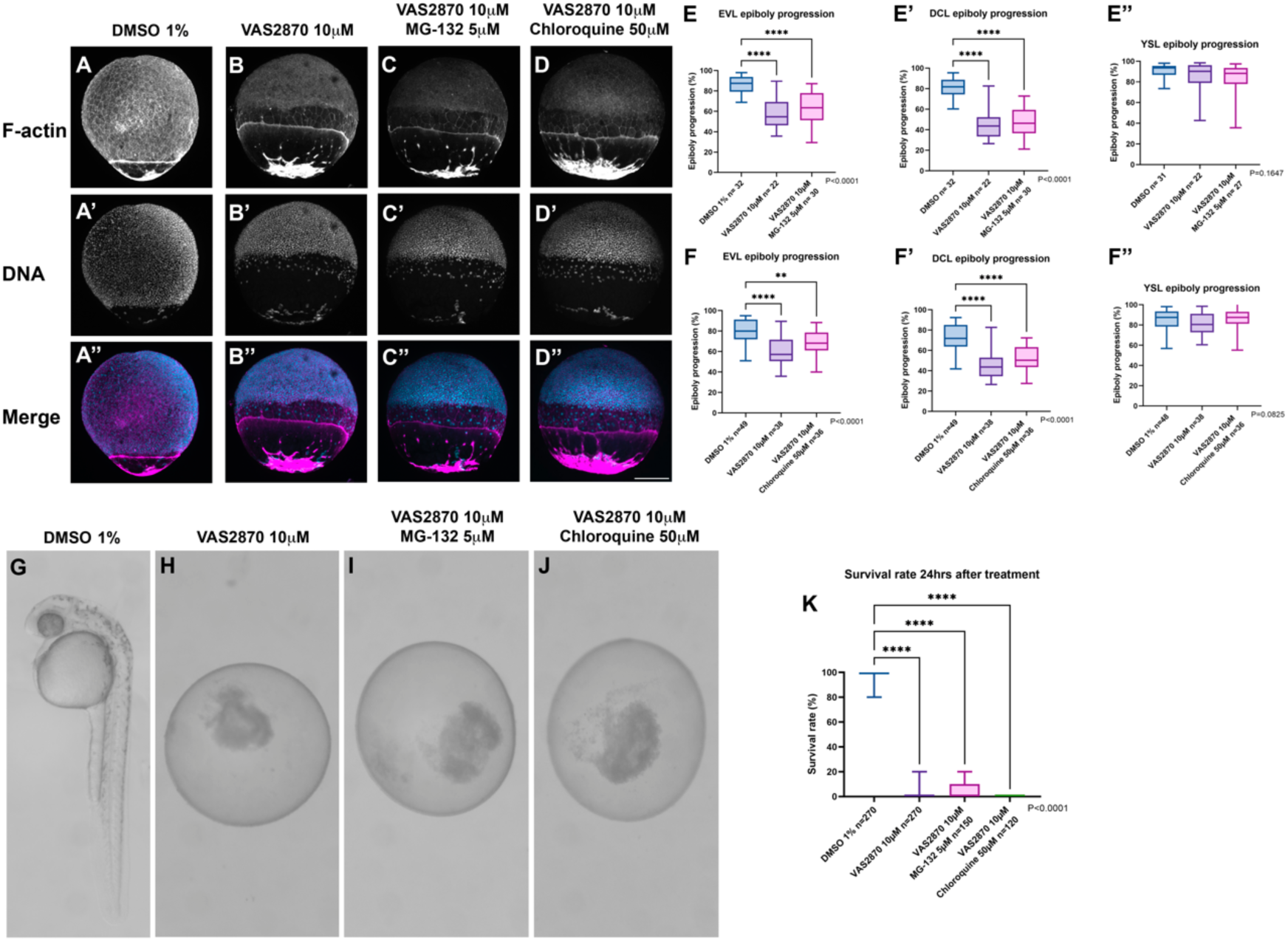
Inhibition of protein degradation systems fail to restore epiboly delay and embryo lethality elicited by Nox activity down regulation. (A-D’’) All treatment groups started with embryos at oblong-sphere stage and the experiment stopped when control treatment embryos reached around 80% epiboly. Embryos were then fixed, stained with DAPI and phalloidin-Alexa and visualized in a spinning disk confocal microscope. (A- A”) Control embryos were incubated with vehicle (1% DMSO). (B-B”) Embryos treated with 10 µM VAS2870 show a delayed epiboly progression. (C-C”) Inhibition of proteasomal protein degradation with 5 µM MG-132 did not rescue the epiboly delay caused by Nox inhibition. (D-D”) Inhibition of lysosomal degradation with 50 µM Chloroquine did not rescue the epiboly delay after treatment with the Nox general inhibitor VAS2870. Scale bar, 200 mm. (E-F”) Enveloping layer (EVL), Deep cell Layer (DCL) and Yolk Syncytial Layer (YSL) epiboly progression quantification in the different treatment groups. Values represent median and interquartile range; whiskers show minimum and maximum values. \**p* < 0.01 by Kruskal-Wallis one-way analysis of variance (ANOVA) with Dunns’s post hoc analysis (*n*, indicated in the graph). N=3 experimental repetitions. (G-H) Inhibition of protein degradation fails to rescue embryo survival. All treatment groups started with embryos at oblong-sphere stage, embryo survival and morphology were analyzed 24hrs after pharmacological treatments (embryos around prim-16/31 hpf). (G) Embryos treated in control media (1% DMSO). (H) Embryos treated with 10 µM VAS2870. (I) Embryos treated with VAS2870 and 5 µM MG-132. (J) Embryos treated with VAS2870 and 50 µM Chloroquine. Scale bar, 300 µm. (K) Quantification of embryo survival 24hrs post treatment with VAS2870 simultaneously with MG-132 and Chloroquine. Values represent median and interquartile range; whiskers show minimum and maximum values. \**p* < 0.01 Kruskal-Wallis one-way analysis of variance (ANOVA) with Dunns’s post hoc analysis (*n*, indicated in the graph). N=3 experimental repetitions.

The simultaneous inhibition of Nox activity and protein degradation with MG-132, chloroquine or H4ClN had no effect on embryo survival, since no embryos remained alive 24 hrs after treatment (Figure 2G to K, and Supplementary Figure 2F and J). We also carried out a survival time curve assays to determine the developmental stage at death after treatment with the inhibitors. The survival rate of embryos treated simultaneously with VAS2870 and MG-132, chloroquine or H4ClN was monitored from the sphere stage up to 24 hpf (Supplementary Figure 5). Cotreatment with VAS2870 and MG-132 or H4ClN further decreased embryo survival when compared with that with VAS2870 at various early stages. This decrease in the survival rate was statistically significant. In contrast, treatment with VAS2870 and chloroquine significantly improved embryo survival between the 70 % epiboly and tailbud stages, delaying embryo death only during this period of development, since no live embryos were observed after 24 hpf of incubation in either treatment group.

Taken together, these results indicate that Nox-derived ROS, in conjunction with dynamin-dependent endocytosis, are required for epiboly progression, embryo survival and development and that their activities are independent of protein degradation.

### Dynasore treatment rescues the effects on E-cadherin and the actin cytoskeleton caused by Nox inhibition

Next, we explored whether the compensatory relationship between Nox inhibition by VAS2870 and endocytosis inhibition by dynasore treatment also compensates for the effects on E-cadherin localization and the actin cytoskeleton, which were previously shown to be affected by VAS2870 treatment (Mendieta-Serrano et al., 2019).

E-cadherin at the EVL cell margins in control embryos exhibited a continuous and intense signal. Additionally, EVL cells exhibited numerous intracellular vesicles that were positive for E-cad (Figure 3A). In contrast, in VAS2870-treated embryos, these signals are depleted, both at the cell margins and in the number of positive vesicles (Figure 3B, G and I). To further characterize this phenomenon, we asked whether endocytosis inhibition with dynasore rescues the effects on E-cad distribution in VAS2870-treated embryos. We found that dynasore treatment restored both the E-cad localization pattern at the EVL cell margins and the number of positive vesicles in VAS2870-treated embryos (Figure 3C, G and I). In contrast, compared with control embryos, DCL cells are not significantly affected in E-cad localization at the cell margins in VAS2870-treated embryos or those subjected to simultaneous treatment with dynasore, (Figure 3D to F, and G’).

**Figure 3.**
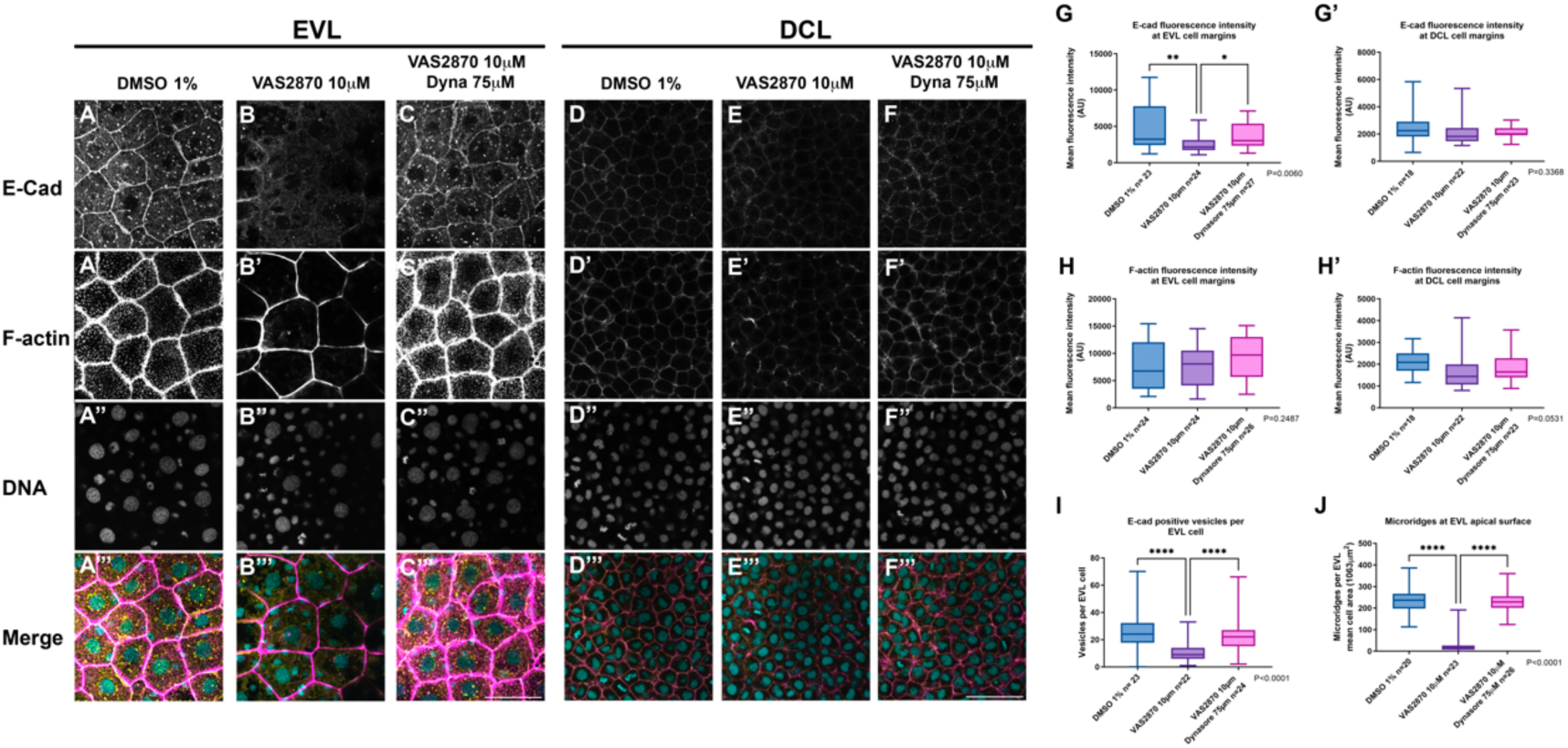
Dynamin activity inhibition by dynasore treatment restores E-cadherin and F-actin localization patterns in zebrafish embryos after Nox activity down regulation. All treatment groups started with embryos at oblong-sphere stage and the experiment stopped when control treatment embryos reached around 65% epiboly. Embryos were then fixed, stained and visualized in a spinning disk confocal microscope. All images were obtained by spinning disk confocal microscopy. E-cad was visualized by immunolocalization in whole embryos in DMSO 1% treated EVL cells (A) and DCL cells (D), 10 µM VAS2870 treatment (B, E) and simultaneous treatment with VAS2870 and 75 µM dynasore (C, F). Filamentous actin (F-actin) was visualized by Phalloidin Alexa Fluor 488 staining of whole embryos in control (A’, D’), 10 µM VAS2870 treatment (B’, E’) and simultaneous treatment with VAS2870 and 75 µM dynasore (C’, F’). Nuclei were visualized in whole embryos by staining DNA with DAPI (A’’-F’’). Images show merged E-cad, F-actin and DNA channels (A’’’-F’’’). Scale bar, 50 µm. E-cadherin fluorescence intensity was measured at EVL (G) and at the DCL (G’) cell margins and compared among treatment groups. Cortical F-actin signal was quantified at EVL (H) and DCL cell margins (H’). The number of vesicles was quantified in the EVL cells in the different treatment groups (I). Microridges per EVL cell were quantified in the different treatment groups (J). Values represent median and interquartile range; whiskers show minimum and maximum values. \**p* < 0.01 by nonparametric one-way analysis of variance (ANOVA) Kruskal Wallis with Dunns’s post hoc analysis (*n*, indicated in the graph). N=3 experimental repetitions. Enveloping layer (EVL). Deep cell layer (DCL).

In our previous report, we showed that in addition to disruption of E-cad localization, the actin cytoskeleton was also affected in EVL cells by VAS2870 exposure (Mendieta-Serrano et al., 2019). On the basis of these findings, we wanted to determine whether inhibiting the early stages of endocytosis could also rescue the effects of VAS2870 treatment on the actin cytoskeleton. We found that the mean fluorescence intensity of the cortical actin network (F-actin) at the EVL (Figure 3A’ and B’) and at the DCL (Figure 3D’ and E’) was not significantly affected in the VAS2870-treated group (Figure 3H and H’ respectively). In contrast, the number of F-actin-rich apical protrusions on EVL cells that correspond to the precursors of microridges (Pinto et al., 2019; Inaba et al., 2020) significantly decreased in VAS2870-treated embryos (Figure 3A’, B’ and J), but these protrusions were recovered on the apical face of the cells by dynasore exposure (Figure 3C’ and J). These results indicate that both E-cad and the actin cytoskeleton are dynamically regulated by endocytosis and that this regulation responds to changes in the availability of Nox-derived ROS.

### Downregulation of Nox-derived ROS increases ZO-1 signal at EVL margins

Adhesion between EVL cells is dependent not only on E-cad adhesion complexes but also on tight junctions. Zonula occludens 1 (ZO-1) is a protein that functions as a scaffold in tight junction assembly, binding to transmembrane occludin and claudins and associating them with the actin cytoskeleton (McNeil et al., 2006). ZO-1 is a cytoplasmic protein that is a member of the membrane-associated guanylate kinase (MAGUK) family of proteins (Stevenson et al., 1986). The localization dynamics of ZO-1, similar to those of E-cad, can be influenced by the endocytosis of tight junction complexes, which are trafficked by clathrin-mediated pathways (Schwarz et al., 2009; Ikari et al., 2011; Farquhar et al., 2012). We tested the effect of Nox inhibition on the localization pattern of ZO-1 at EVL cell margins to evaluate whether the changes in adhesion protein localization were specific to E-cad or if they involved other proteins. In control embryos (Figure 4A), ZO-1 shows a strong membrane signal at the EVL cell margins, as previously reported by others (Song et al., 2013; Lepage et al., 2014), as well as a granulated pattern in the nucleus, because the protein has two nuclear localization domains (Gonzalez-Mariscal et al., 1999). Interestingly, after Nox inhibition, the ZO-1 signal increased significantly at the EVL cell margins and showed no patches or discontinuous patterns (Figure 4B and E). This finding indicates that, unlike adherens junctions, tight junctions are not disturbed by the inhibition of Nox activity. In contrast, tight junctions would appear to increase, possibly as a compensatory mechanism for the loss of E-cadherin function.

**Figure 4.**
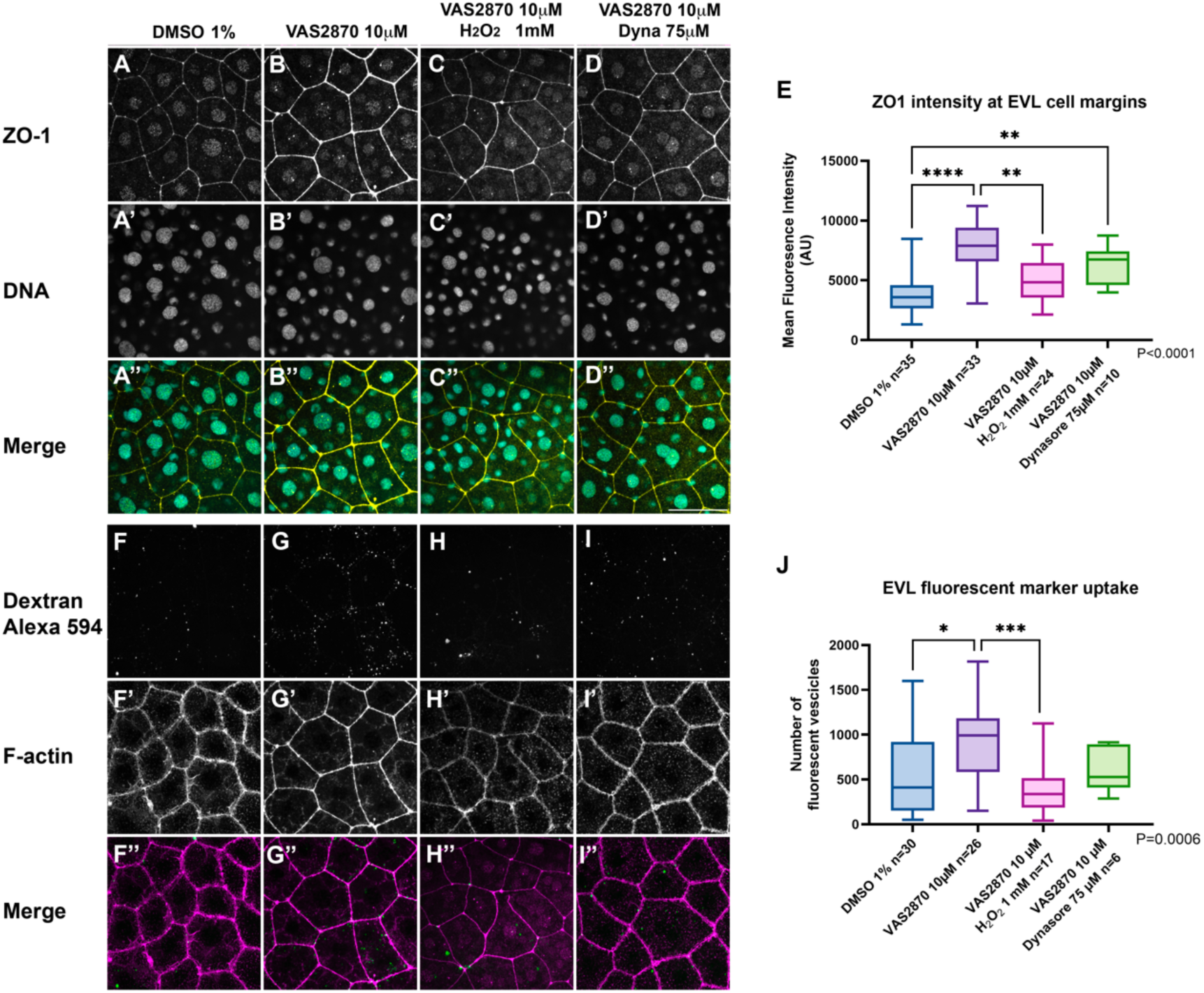
Nox inhibition increases ZO-1 signal and endocytosis at EVL cell margins, while H_2_O_2_ and Dynasore treatment restores ZO-1 pattern endocytosis. All treatment groups started with embryos at oblong-sphere stage and the experiment stopped 3 hrs after the start of the treatment (around 60% epiboly). Embryos were then fixed, stained and visualized in a spinning disk confocal microscope. ZO-1 was visualized by immunolocalization in whole embryos in DMSO 1% treated EVL cells (A), 10 µM VAS2870 treatment (B), simultaneous treatment with VAS2870 and H_2_O_2_ 1 mM (C), and simultaneous treatment with VAS2870 and 75 µM dynasore (D). Nuclei were visualized in whole embryos by staining DNA with DAPI (A’-D’). Images show merged ZO-1 and DNA channels (A’’- D’’). Scale bar, 50 µm. Quantification of ZO-1 signal at EVL cell margins (E). Values represent median and interquartile range; whiskers show minimum and maximum values. \**p* < 0.01 by nonparametric one-way analysis of variance (ANOVA) Kruskal Wallis with Dunns’s post hoc analysis (*n*, indicated in the graph). Enveloping layer (EVL). N=3 experimental repetitions. (F-J) Increased fluid phase endocytosis elicited by Nox activity down regulation is brought back down by H_2_O_2_ treatment. All images were obtained by spinning disk confocal microscopy. Fluid phase endocytosis was visualized in EVL cells by adding fluorescent dextran to the embryo culture media. Dextran 594 was visualized in control embryos (DMSO 1%) (F), 10 µM VAS2870 treatment (G), simultaneous treatment with VAS2870 and 1 mM H_2_O_2_ (H), and simultaneous treatment with VAS2870 and 75 µM dynasore (I). Filamentous actin (F-actin) was visualized by Phalloidin Alexa Fluor 488 staining of whole embryos in control (F’-I’). Images show merged Dextran 594 and F-actin channels (F’’-I’’). Scale bar, 50 µm. Quantification of the number of fluorescent vesicles found in EVL cells (J). Values represent median and interquartile range; whiskers show minimum and maximum values. \**p* < 0.01 by nonparametric one-way analysis of variance (ANOVA) Kruskal Wallis with Dunns’s post hoc analysis. (*n*, indicated in the graph. N=3 experimental repetitions.

When embryos were treated simultaneously with VAS2870 and 1 mM H_2_O_2_, the intensity of the ZO-1 signal decreased and returned to control levels (Figure 4C and E), as did embryos treated with VAS2870 and the dynamin inhibitor dynasore (Figure 4D and E). These results indicate that the decrease in Nox-derived ROS causes an increase in tight junction components at the EVL, and that this increase is regulated at least in part by endocytosis.

### Nox activity downregulation increases fluorescent fluid-phase marker uptake in EVL cells

Since the regulation of E-cad endosomal trafficking is important for blastoderm cell movement and epiboly progression (Song et al., 2013) and since the epiboly delay caused by VAS2870 decreases E-cad localization or abundance at the EVL cell margins (Mendieta-Serrano et al., 2019), we hypothesized that the ROS produced by NADPH oxidases negatively regulate endocytosis. Accordingly, we propose that the changes in the pattern of E-cad localization or abundance observed in this work (Figure 3) may be indicative of membrane protein turnover variations, which are affected by the endocytic flow and recycling. To directly probe into the endocytic rate, we analyzed the uptake of the fluorescent fluid-phase marker dextran Alexa 594 (Sonal et al., 2014). Dechorionated embryos at the oblong-sphere stage were incubated for 2 hours in ERM media containing the inhibitors and the fluid-phase marker, and after this treatment, the embryos were fixed and costained with DAPI and Alexa Fluor 488 phalloidin. Visualization of the fluorescent endosomes found in EVL cells, despite showing a low, dispersed and variable signal (Figure 4F to I), revealed visible differences between the treatment groups. We focused on analyzing endocytosis in EVL cells since significant changes in E-cad abundance were detected in VAS2870-treated embryos. In all the treatment groups, the EVL cells exhibited small fluorescent structures of slightly different sizes and signal intensities, which we consider to be presumptive endocytic vesicles (white arrows in Figure 4F to I). When embryos were treated with VAS2870, the number of vesicles loaded with the fluorescent dye was significantly greater in the EVL cells-treated group than in the control treatment group (Figure 4G and J). The localization pattern of ZO-1 at the EVL cells margins in VAS2870-treated embryos (Figure 4B) rules out the possibility that the fluorescent dextran signal observed at the EVL was caused by permeability changes at the cell junctions due to a decrease in adhesion. Therefore, the increased number of fluorescent vesicles indicates an increased rate of fluid-phase endocytosis at the EVL apical surface and not by fluorescent dye seeping in between the cell margins.

We tested our proposed Nox modulation of endocytosis by adding H_2_O_2_ to VAS2870- treated embryos or by the simultaneous inhibition of Nox and dynamin activity. H_2_O_2_ is a ROS that is presumably downregulated in embryos exposed to VAS2870 and is capable of rescuing its effects on epiboly, cell motility, embryo survival, and changes in E-cad localization and cytoskeleton stability (Mendieta-Serrano et al., 2019).

Cotreatment with VAS2070 and 1 mM H_2_O_2_ decreased the intake of fluorescent dye, restoring the number of fluorescent vesicles to control levels (Figure 4H and J). The same effect was achieved by simultaneous treatment with VAS2870 and dynasore (Figure 4I and J), although the magnitude of the rescue observed was greater with H_2_O_2_. These results strongly suggest that H_2_O_2_ produced by Nox activity, similar to dynasore activity, acts as a negative regulator of endocytosis. Therefore, while the absence of H_2_O_2_ caused by the loss of Nox activity elicits an increase in fluid-phase endocytosis, the addition of exogenous H_2_O_2_ prevents this surge and restores endocytosis to control levels.

Strikingly, treating embryos with only a high concentration of H_2_O_2_ (2.5 mM), significantly decreased the number of fluorescent vesicles compared with that of control embryos (Supplementary figure 7), further supporting the role of Nox-derived ROS, particularly H_2_O_2_, as negative regulators of the endocytic rate and fluid-phase endocytosis.

### Nox-derived ROS regulate the localization of the endocytosis-trafficking machinery

Since Nox-derived ROS participate in the modulation of endocytosis, we wanted to address whether any of the endocytic or vesicular transport machinery components are affected by VAS2870 treatment.

Although there are different E-cadherin internalization mechanisms, we focused our attention on clathrin-dependent endocytosis, since it seems to be the internalization mechanism most commonly used in mammalian cells, the cell model best studied to date (Cadwell et al., 2016). First, we aimed to analyze the localization patterns of three components of the clathrin-dependent pathway known to be expressed in early zebrafish embryos, clathrin (Feng et al., 2002; Eno et al., 2016), dynamin 2 (Dnm2) (Eno et al., 2016) and a protein important for endosome recycling, Rab11 (Zhang et al., 2019), using 3 different antibodies previously reported to produce positive signals in early developing zebrafish embryos (Eno et al., 2016).

Since the signals corresponding to Dnm2 and clathrin were highly variable between the samples and the results were not very consistent (not shown), we focused on the immunolocalization pattern of Rab11, which showed a reproducible and distinct localization pattern among all the replicas in all the treatment groups. In control embryos, Rab11 has a dispersed intracellular distribution and an abundant signal at the plasma membrane and its vicinity (Figure 5A). After Nox inhibition with VAS2870 treatment, Rab11 fluorescent signal at the membrane significantly decreased when compared to control embryos (Figure 5B and E). These findings indicate that Nox inhibition hinders the transport of Rab11-associated vesicles toward the plasma membrane, impeding cargo recycling. Notably, simultaneous treatment with VAS2870 and H_2_O_2_ significantly restored the Rab11 signal at the plasma membrane (Figure 5C and E). Compared with VAS2870 treatment, cotreatment with dynasore also significantly increased the Rab11 signal at the EVL cell membrane; however, compared with that in the control group, the Rab11 signal in the EVL cell membrane was not completely restored. (Figure 5D and E). There were no differences in the cytoplasmic Rab11 signal between the treatment groups (Figure 5A to D and F).

**Figure 5.**
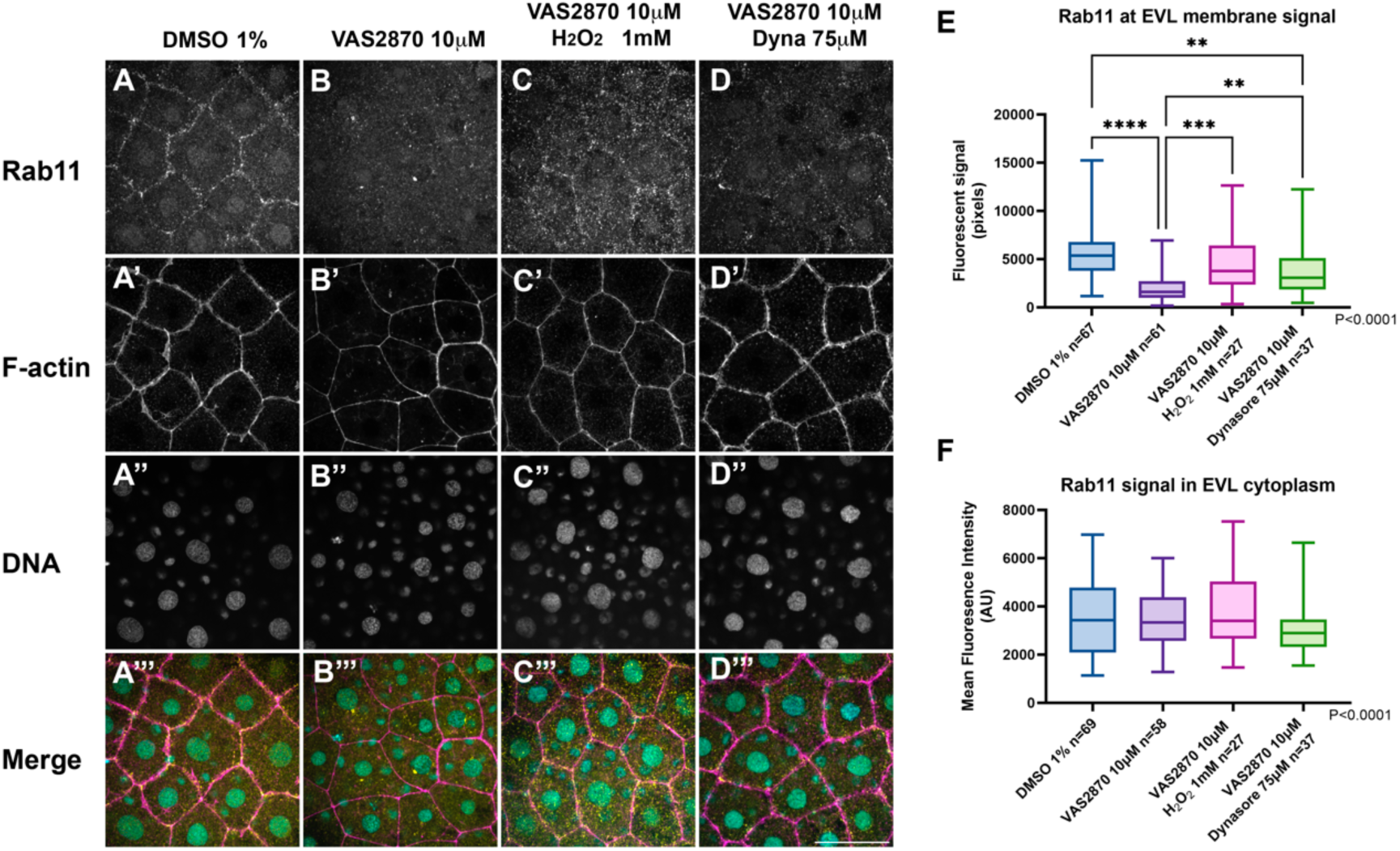
Loss of Rab11 signal at the plasma membrane in EVL cells caused by Nox activity down regulation is prevented by H_2_O_2_ treatment. All treatment groups started with embryos at oblong-sphere stage and the experiment stopped when control treatment embryos reached around 65% epiboly. All images were obtained by spinning disk confocal microscopy. Rab11 was visualized in control embryos (DMSO 1%) (A), 10 µM VAS2870 treatment (B), simultaneous treatment with VAS2870 and 1 mM H_2_O_2_ (C), and simultaneous treatment with VAS2870 and 75 µM dynasore (D). Filamentous actin (F- actin) was visualized by Phalloidin Alexa Fluor 488 staining (A’-D’). Nuclei were visualized by staining DNA with DAPI (A’’-D’’). Images show merged Rab11, F-actin and DNA channels (A’’-D’’). Scale bar, 50 µm. H_2_O_2_ treatment significantly increases Rab11 signal at the EVL cell margins after VAS2870 treatment, while dynamin activity inhibition by dynasore treatment also increased Rab11 signal but fails to reach control levels (E). Treatments have no effects on cytoplasmic Rab11 signal levels (F). Values represent median and interquartile range; whiskers show minimum and maximum values. \**p* < 0.01 by nonparametric one-way analysis of variance (ANOVA) Kruskal Wallis with Dunns’s post hoc analysis. (*n*, indicated in the graph). N=3 experimental repetitions.

These results indicate that downregulation of Nox-derived ROS by the treatment with VAS2870 affects the abundance and pattern of a Rab GTPase associated with recycling endosomes back to the plasma membrane. Although the inhibition of endocytosis via dynasore treatment is capable of partially rescuing Rab11 localization at the plasma membrane, only supplementation with exogenous H_2_O_2_ effectively restores the distribution to an equivalent control pattern, indicating that Nox-derived ROS and dynasore work at different levels.

Our results suggest that Nox activity and its products can influence the distribution of proteins important for the endocytic trafficking machinery and that H_2_O_2_ participates in the regulation of protein recycling back to the plasma membrane in a Rab11-dependent manner.

## DISCUSSION

### Nox derived ROS act as a negative regulator of endocytosis during early development in zebrafish

Endocytosis is an essential process that involves the internalization of different molecules present in the extracellular fluid-phase along with proteins and lipids contained in the cell membrane. Endocytosis is involved not only in basic cell physiological functions such as nutrient internalization but also affect diverse macromolecules present at the plasma membrane. Endocytosis internalizes ligands and receptors that enter the cytoplasm and are transported in the cells by vesicular trafficking. Therefore, endocytosis plays key roles in the regulation of signaling, cell adhesion and migration, among many other processes important for the development of multicellular organisms and in adult life homeostasis (Doherty and McMahon, 2009; Kaksonen and Roux, 2018). For cell motility, coordination during epiboly endocytosis appears to be crucial. In pioneering work with the killifish *Fundulus heteroclitus*, Betchaku and Trinkaus described intense endocytosis at the external yolk syncytial layer (E-YSL) just in front or vegetally to the EVL (Betchaku and Trinkaus, 1986). Since then, it has been proposed that endocytosis at the E-YSL, most likely micropinocytosis (Marsal et al., 2021), promotes epiboly progression of the blastoderm (Betchaku and Trinkaus, 1986; Solnica-Krezel and Driever, 1994). However, endocytosis at the blastoderm also seems to contribute to epiboly progression. Global inhibition of dynamin-dependent endocytosis in whole embryos delays epiboly, but when dynamin 2 (Dnm2) activity is disrupted by restricted expression of a dominant-negative Dnm2 in yolk cell, blastoderm and yolk syncytium nuclei epiboly is not significantly affected (Lepage et al., 2014). However, different results were obtained when the small GTPase Rab5ab was downregulated by morpholino injection into the yolk cell after the midblastula transition. Here, the knockdown of Rab5ab decreased E-YSL endocytosis and delayed epiboly (Marsal et al., 2021). These contrasting results suggest that endocytosis at both the blastoderm and the E-YSL might contribute to epiboly. In the present work, we found that Nox-derived ROS, more likely H_2_O_2_, contribute together with dynamin-dependent endocytosis to promote blastoderm epiboly progression. First, we confirmed that the general pharmacological inhibition of Nox with VAS2870 in embryos at the sphere stage delays epiboly, downregulates E-cad in EVL cells and affects embryo development and survival. All these effects are rescued by dynasore (Figures 1 and 3). We propose that epiboly requires the modulation of endocytosis, and this modulation is achieved by the negative effect of Nox-derived ROS. When these ROS are not produced, the inhibition of endocytosis by dynasore allows the proper advancement of epiboly (Figure 1).

### Nox catalytic activity and H_2_O_2_ decreased fluid-phase endocytosis

Importantly, we confirmed that Nox inhibition increases fluorescent fluid-phase endocytosis in EVL cells, whereas either H_2_O_2_ or dynasore cotreatments decreases dextran-Alexa internalization, thereby restoring fluid-phase endocytosis to a control treatment equivalent level (Figure 4). These results show that H_2_O_2_ is involved in the modulation of endocytosis. Moreover other groups have reported that H_2_O_2_ treatment inhibits epidermal growth factor receptor internalization in a fibroblast line *in vitro* (De Wit et al., 2000; De Wit et al., 2001), suggesting that H_2_O_2_ participation in endocytosis modulation is not specific to zebrafish embryonic cells.

In the present work, we focused on characterizing the effects on the EVL apical surface endocytosis; however, it will be important to characterize in detail the effects on the endocytosis on the basolateral side of the EVL cells, the side that interacts with the deep cells (DCs) and contributes to the spread of DC epiboly.

Nevertheless, there is a potential alternative cause for these results. The significant increase in fluorescent fluid-phase endocytosis markers in EVL cells in VAS2870-treated embryos could be due not only to an increased endocytosis but also to compromised EVL barrier function. However, we found that the abundance of the tight junction scaffold protein ZO- 1 significantly increased at the EVL cells margins in VAS29870-treated embryos (Figure 4). This result indicates that tight junctions are not compromised in these treated embryos and potentially also shows that a compensatory process is stimulated due to the decrease in E-cadherin at the EVL membrane.

### Nox-derived ROS, particularly H_2_O_2_, promote Rab11 localization at the plasma membrane

We also found that Rab11, which is a well-known monomeric small GTPase involved in the regulatory system of recycling endosome trafficking (Ullrich et al., 1996), is affected by the inhibition of Nox. Rab11 localization and intensity are affected in VAS2870-treated embryos (Figure), indicating that endosome recycling is potentially disrupted. Other small GTPases of the Rab11 subfamily that are known for their role in endosome recycling, such as Rab25 (Casanova et al., 1999), have been shown to be required for epiboly progression (Willoughby et al., 2021). In our experiments, we found that H_2_O_2_ or dynamin inhibition revered the effects on Rab11. These results might explain the decrease in the E-cad signal at the EVL cell margins in VAS2870-treated embryos, potentially due to the decrease in E-cad recycling back to the plasma membrane. However, we do not yet have a reasonable explanation for the exact mechanism behind the overall decline in the E-cad signal both at the plasma membrane and at intracellular vesicles in VAS2870-exposed embryos. We hypothesized that protein degradation inhibition of either the proteasome system or the lysosome system would rescue these embryos. However, in these cotreatment schemes, we were unable to find evidence of any rescue for epiboly progression, general development or survival. These results suggest that an increase in protein degradation is not the main cause of the developmental effects observed after decreasing ROS formation.

Some of the observed changes in the abundance and pattern of Rab11 proteins could be mediated by the effects on the cytoskeleton. Nox inhibition altered F-actin (Figure 3) and the microtubule cytoskeleton, as previously reported (Mendieta-Serrano et al., 2019). Some particular Rab11-family interacting proteins (Rab11-FIPs) mediate the interaction of Rab11 with motor proteins such as dynein, myosin V and kinesin, and regulate their interaction with microtubules to mediate their movement and localization (Horgan et al., 2010). Our results suggest that Nox ROS possibly participate in the regulation of Rab11 localization and stability, which are important for endosome recycling through their effects on the actin or tubulin cytoskeleton.

Since our results indicate that Nox-derived ROS contribute to the regulation of endocytosis and vesicular trafficking at different levels, this led us to ask how this process is related to the regulation of epiboly? For epiboly to successfully take place, cell mechanical forces and biochemical signals, together with diverse cellular processes (Schwayer et al., 2019), such as endocytosis, are required to act in concert among the different cells that constitute the developing embryo. The prevailing view is that the external yolk syncytial layer (E-YSL) is the main motor of epiboly that generates the mechanical force necessary to pull the blastoderm toward the vegetal pole, owing to the strong attachment of the EVL margin to the E-YSL (Trinkaus, 1984; Lepage and Bruce, 2010; Lepage et al., 2014). In fact, it was shown in *Fundulus heteroclitus* that the YSL is capable of undergoing epiboly independently when the blastoderm is removed or when the EVL attachment to the E-YSL is severed; furthermore, these experimental manipulations accelerate epiboly progression, indicating that E-YSL dragging of the blastoderm retards epiboly (Trinkaus, 1984). The EVL vegetal margin is strongly attached to the YSL through connecting tight junctions that are fundamental for the traction force of the YSL upon the EVL (Koppen et al., 2006). In addition, actomyosin rings located at the edge of EVL and at the YSL are also required for driving the spread of EVL over the yolk cell through ring contractions and a flow-friction mechanism (Behrndt et al., 2012). Although the EVL epiboly spread appears to be a partially passive process (Bruce and Heisenberg, 2020), active participation of the EVL is also required since these cells undergo major structural rearrangements, such as characteristic cell flattening, by decreasing their height and dramatically enlarging their apical surface (Campinho et al., 2013). In *Fundulus heteroclitus*, dramatic rearrangements of EVL cells during epiboly require intense membrane turnover by endocytosis and exocytosis. which mainly occur at the cell margins, and that membrane turnover is increased during epiboly via increased mechanical tension (Fink and Cooper, 1996). The entire periphery of EVL cells is mechanically coupled by adherent and tight junctions, and dynamin is required for the localization and stability of both kinds of junctions (Lepage et al., 2014). As indicated by our results, it is tempting to propose that Nox-derived ROS participate in the fine tuning of the rate of endocytosis in EVL cells, a process that impacts different cell junctions and their response to the changes in mechanical tension required for the necessary structural rearrangements that occur at the EVL necessary for normal epiboly.

Overall, our results reveal new roles for Nox-derived ROS in regulating endocytosis and trafficking of plasma membrane components in EVL cells, which are necessary for proper epiboly regulation. Finally, importantly, the participation of H_2_O_2_ in the modulation of endocytosis seems to be evolutionarily conserved. *In vitro* exposure to H_2_O_2_ has previously been shown to reversibly inhibit/delay endocytosis in evolutionarily distant organisms such as yeast (Pereira et al., 2012) and in different human cell types in tissue culture (De Wit et al., 2000; De Wit et al., 2001; Cheng and Vieira, 2006). Exploring the potential evolutionary origins of this interesting regulatory mechanism in more detail is important.

## ACKNOWLEDGEMENTS

We are grateful to the reviewers for their critical comments on the manuscript. We thank Laura Ramirez for technical support and Dulce Pacheco for animal care. Confocal microscopy technical support provided by Arturo Pimentel, and service by Laboratorio Nacional de Microscopía Avanzada, UNAM. We thank Mario Mendieta-Serrano for the support with the F-actin and E-cadherin quantification analysis. We acknowledge the contribution of Francisco J. Mendez-Cruz at the start of the project. This work was supported by grants given to E. S-V from PAPIIT-UNAM, IN212820 and IN227223, H.L. DGAPA-UNAM grant IN206822 and CONAHCyT 128353. A.R.C. acknowledges fellowships 720706 and 924853, and B.R.M. 1314107 from CONAHCyT, in addition to fellowships from PAPIIT-UNAM, project IN212820.

## AUTHOR CONTRIBUTIONS

E.S.-V., and A.R.C. designed the project; A.R.C. performed most of the experiments, the imaging and quantification analysis; B.R.-M performed the rescue experiments with MG- 132, chloroquine and H4ClN; E.S.-V., A.R.C., D.S.P., and H.L. wrote the paper.

---

**Supplementary Figure 1.**
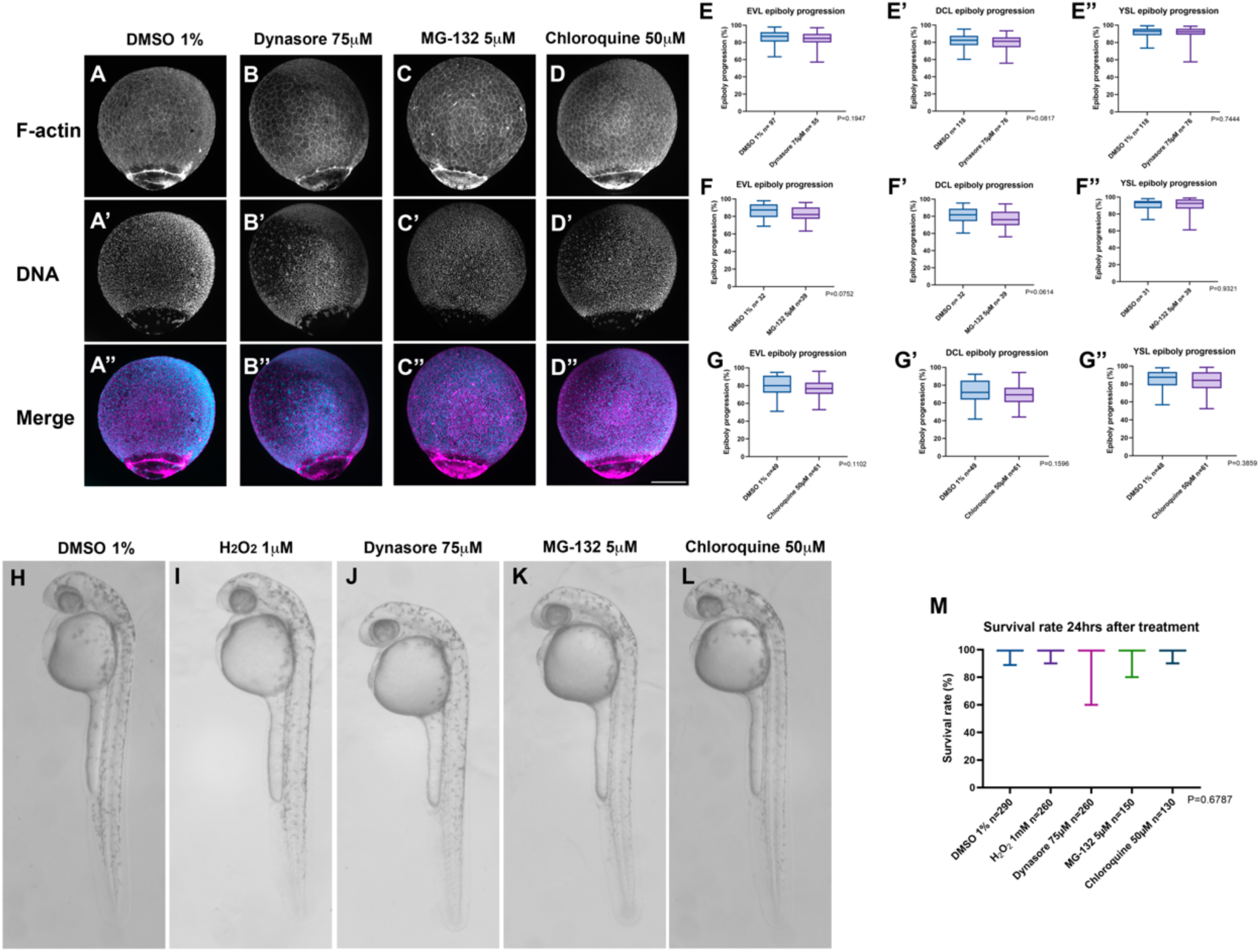
Treatment with Dynasore, MG-132 and Chloroquine added individually to the culture media show no effect on epiboly progression and embryo survival. (A-G”). All treatment groups started with embryos at oblong-sphere stage and the experiment stopped when control treatment embryos reached around 80% epiboly. Embryos were then fixed, stained with DAPI and phalloidin-Alexa and visualized in a spinning disk confocal microscope. (A-A”) Control embryos were incubated with vehicle (1% DMSO). (B-B”) Embryos treated with 75 µM Dynasore (Dyna). Embryos treated with 5 µM MG-132. (C-C”). Embryos treated with 50 µM Chloroquine. Scale bar, 200 µm. (E- G”) Enveloping layer (EVL), Deep cell Layer (DCL) and Yolk Syncytial Layer (YSL) epiboly progression quantification in the different treatment groups. Values represent median and interquartile range; whiskers show minimum and maximum values. Data analyzed with Mann-Whitney or Welch’s t Test. No treatment showed a significant effect. (*n*, indicated in the graph). N=3 experimental repetitions. (H-L) Treatment with 1 mM H_2_O_2_, 75 mM Dynasore, 5 mM MG-132 and 50 mM Chloroquine added individually to the culture media show no effect on epiboly progression. All treatment groups started with embryos at oblong-sphere stage and embryo survival and morphology were analyzed 24h after pharmacological treatments. Scale bar, 300 µm. (M) Quantification of live embryos at 24 hrs post treatment. No treatment showed a significant effect on embryo survival. Values represent median and interquartile range; whiskers show minimum and maximum values. *P*=0.9362 by Kruskal-Wallis one-way analysis of variance (ANOVA) with Dunns’s post hoc analysis (*n*, indicated in the graph). N=3 experimental repetitions.

**Supplementary Figure 2.**
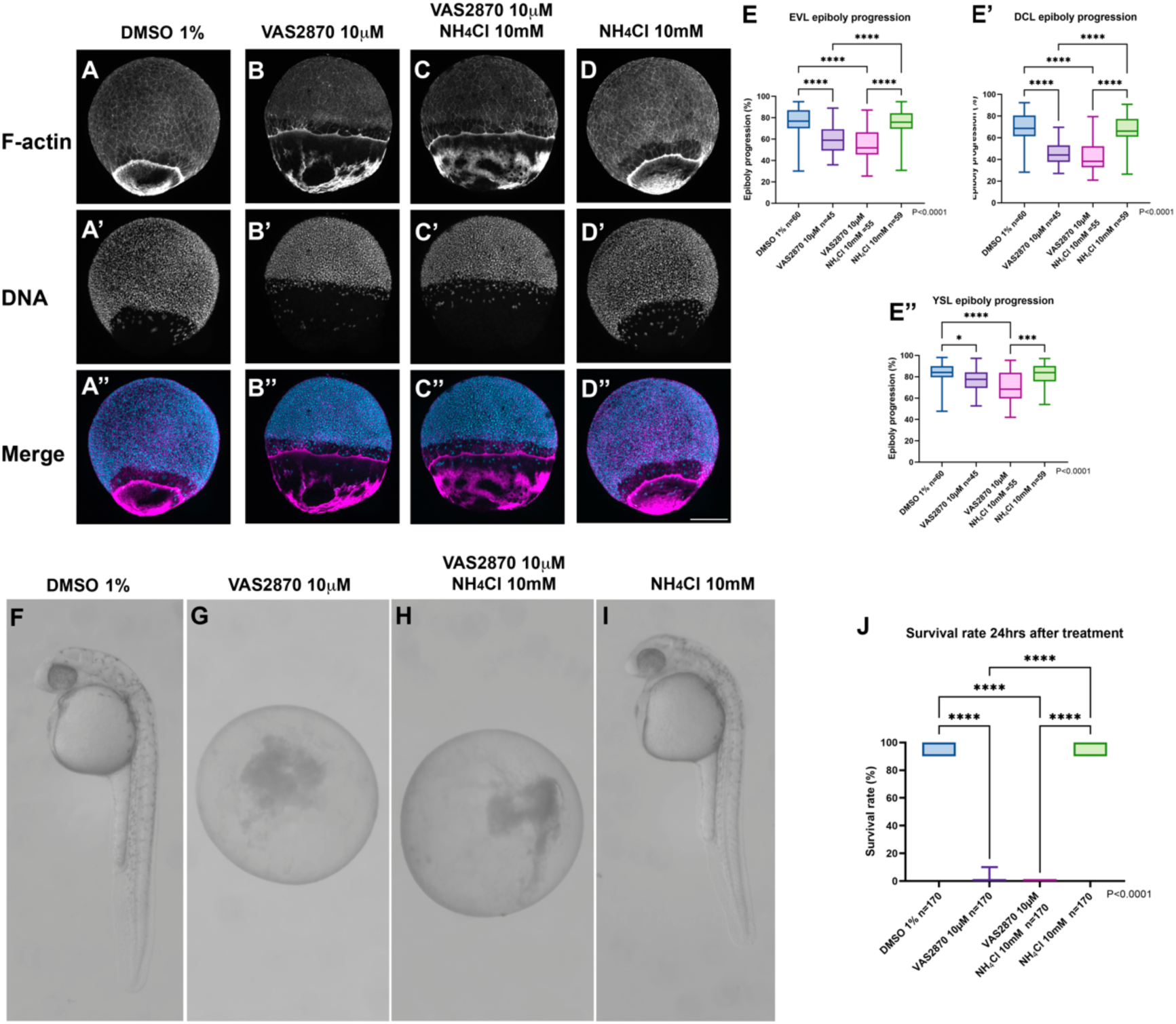
Inhibition of lysosome activity with ammonium chloride (NH4Cl) fails to rescue epiboly progression after Nox inhibition. (A-D”) All treatment groups started with embryos at oblong-sphere stage and the experiment stopped when control treatment embryos reached around 80% epiboly. Embryos were then fixed, stained with DAPI and phalloidin-Alexa and visualized in a spinning disk confocal microscope. (A-A”) Control embryos were incubated with vehicle (1% DMSO). (B- B”) Embryos treated with 10 µM VAS2870 show a delayed epiboly progression. (C-C”) Delayed epiboly progression after treatment with the Nox general inhibitor VAS2870 shows no improvements after treatment with the lysosomal inhibitor 10 mM NH4Cl. (D, D”) Treatment with 10 mM NH4Cl has no effect on epiboly progression when compared to control embryos. Scale bar 200 µm. (E, E”) Quantification of enveloping layer (EVL), deep cell layer (DCL) and yolk syncytial layer (YSL) epiboly progression. Values represent median and interquartile range; whiskers show minimum and maximum values. \**p* < 0.01 by Kruskal-Wallis one-way analysis of variance (ANOVA) with Dunns’s post hoc analysis (*n*, indicated in the graph). N=3 experimental repetitions. (F-I) The simultaneous treatment with 10 mM NH4Cl and VAS2870 shows effect on embryo survival and development. All treatment groups started with embryos at oblong-sphere stage and embryo survival and morphology were analyzed 24h after pharmacological treatments. Scale bar 300 µm. (J) Quantification of live embryos at 24 hrs post treatment. All boxes represent median and interquartile range; whiskers show minimum and maximum values. \**p* < 0.01 by Kruskal-Wallis one-way analysis of variance (ANOVA) with Dunns’s post hoc analysis (*n*, indicated in the graph). N=3 experimental repetitions.

**Supplementary Figure 3.**
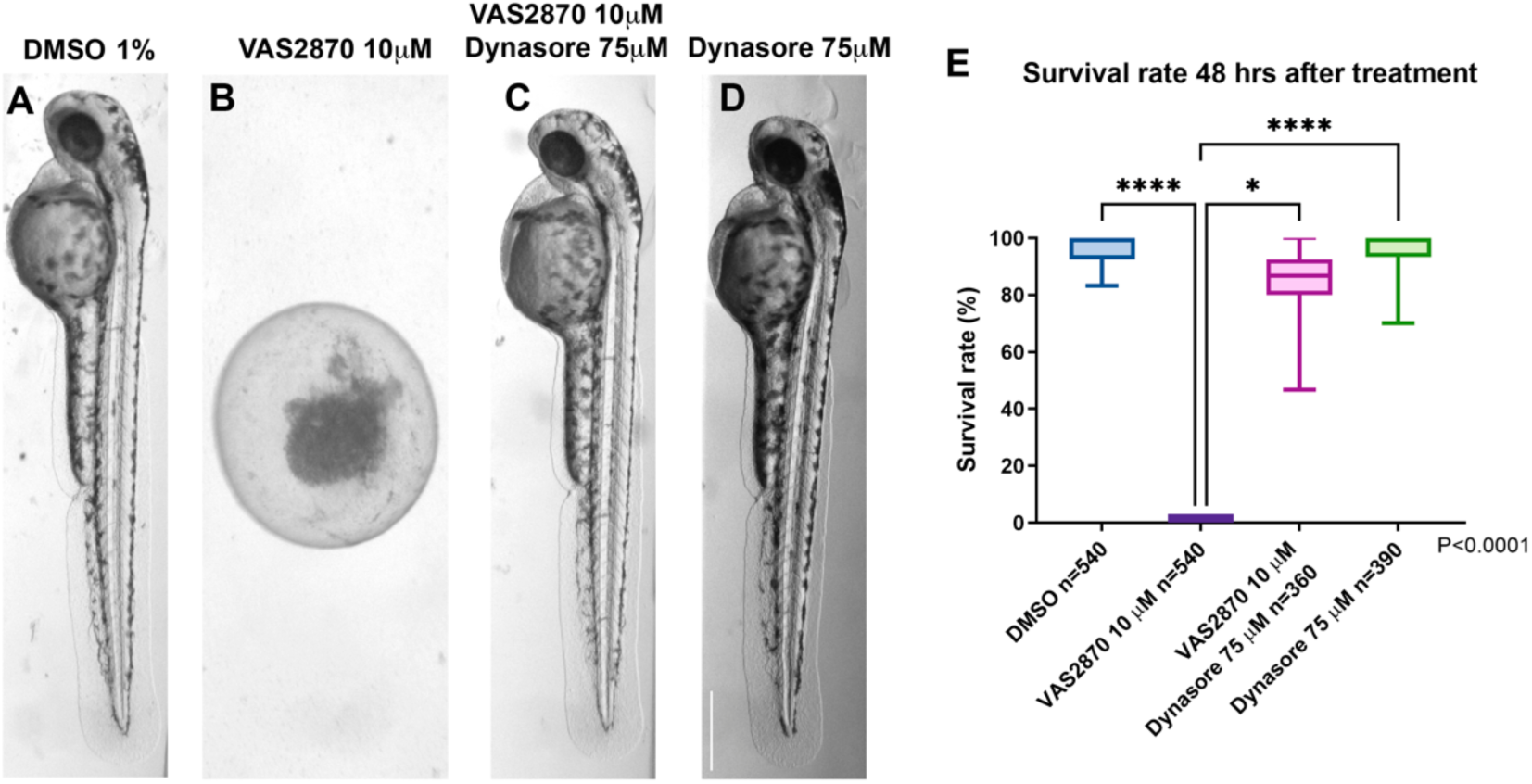
Inhibition of endocytosis rescues embryo survival and development affected by Nox activity downregulation at 48 hpf. All treatment groups started with embryos at oblong-sphere stage, embryo survival and morphology were analyzed 48 hrs after pharmacological treatments. (A) Embryos treated in control media (1% DMSO). (B) Embryos treated with 10 µM VAS2870. (C) Embryos treated with VAS2870 and 75 µM Dynasore simultaneously. (D) Embryos treated with 75 µM Dynasore (E). Scale bar 300 µm. Quantification of embryo survival 48 hrs post treatment. Values represent median and interquartile range; whiskers show minimum and maximum values. \**p* < 0.01 Kruskal-Wallis one-way analysis of variance (ANOVA) with Dunns’s post hoc analysis (*n*, indicated in the graph). N=3 experimental repetitions.

**Supplementary Figure 4.**
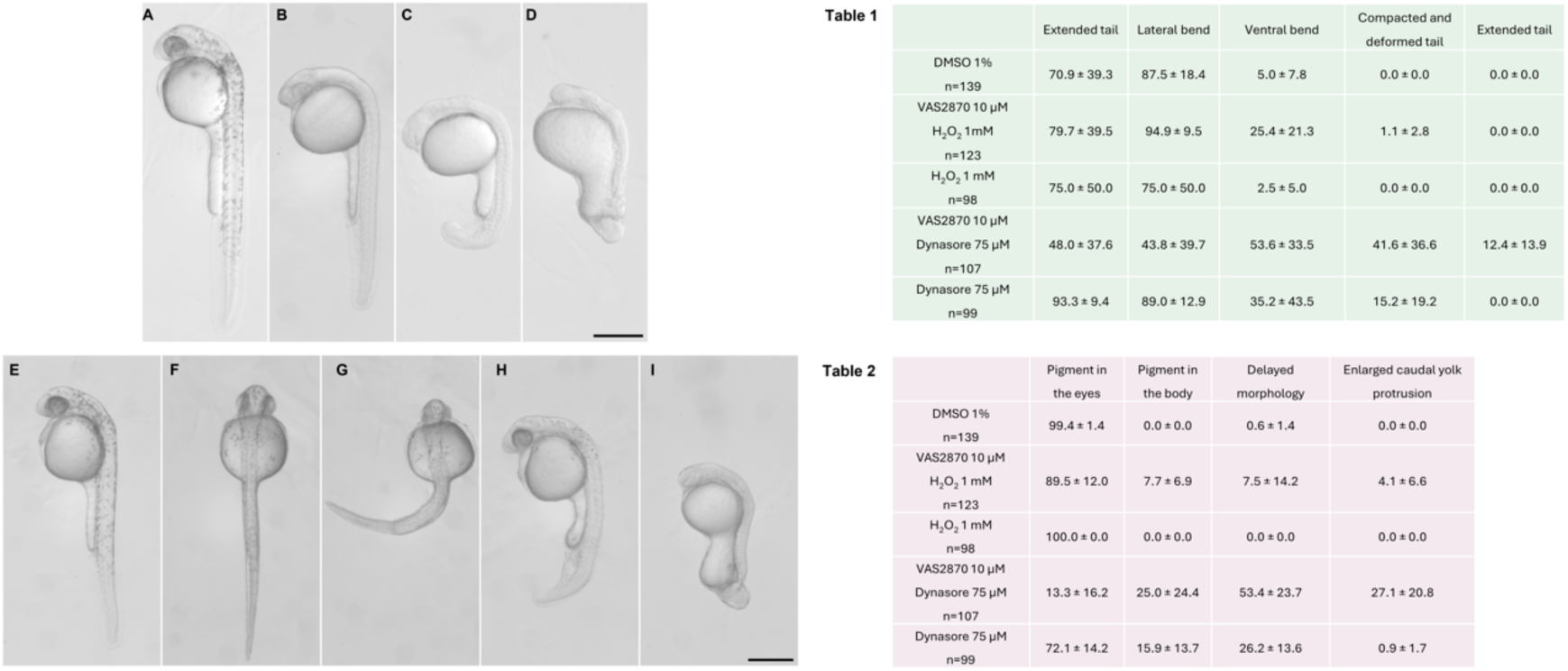
Phenotypes observed in embryos treated simultaneously with Dynasore and VAS2870 display a gradient of both general and caudal defects. (A) Embryo shows a normal pigmentation pattern found in both eyes and body (B). Embryo shows a lack of pigmentation in the eyes and the body. Morphology also seems developmentally delayed. (C) Embryo appears to be significantly delayed in comparison to other littermates and presents an enlarged caudal yolk protrusion. (D) Embryo is severely delayed and deformed, with various tissues and structures affected. Scale bar 300 µm. Table 1 shows the frequency of the observed phenotypes found in the different treatment groups. (E) Embryo shows a normal caudal morphology in a lateral and dorsal (F) view. (G) Embryo presents a lateral caudal bend. (H) Embryo with a ventral bend of the tail. (I) Embryo with a severely deformed tail. Scale bar 300 µm. Table 2 shows the frequency of the observed caudal phenotypes in the different treatment groups.

**Supplementary Figure 5.**
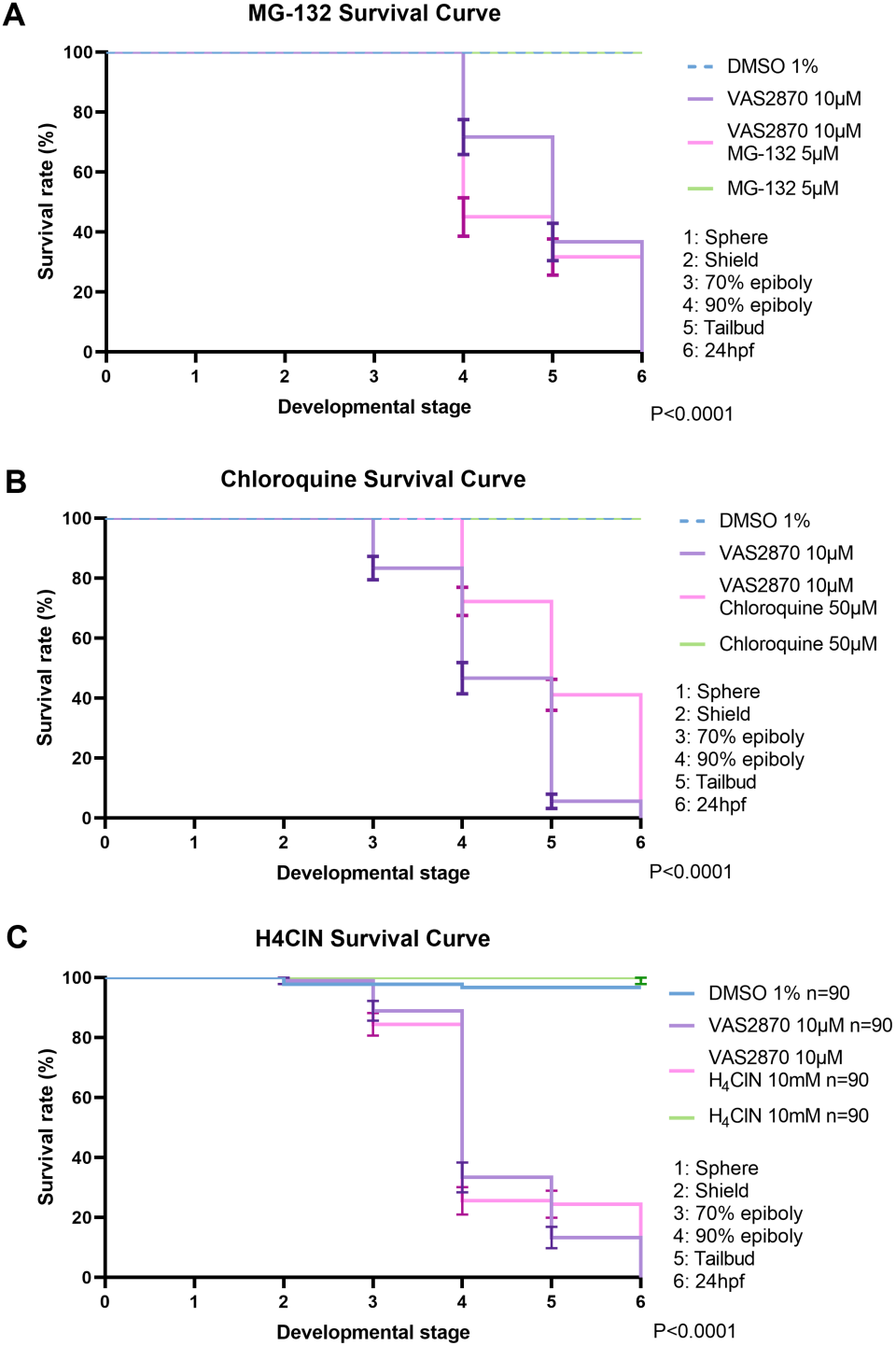
Inhibition of lysosome activity with chloroquine significantly improved embryo survival between the 70 % epiboly and tailbud stages. (A) MG-132 treatment decreases embryo survival between the 70 % epiboly and tailbud stages in embryos cotreated with VAS2870. (B) Chloroquine treatment decreased embryo survival between the 70 % epiboly and tailbud stages in embryos cotreated with VAS2870. (C) Ammonium chloride treatment decreased embryo survival between the 70 % epiboly and tailbud stages in embryos cotreated with VAS2870 treatment

**Supplementary Figure 6.**
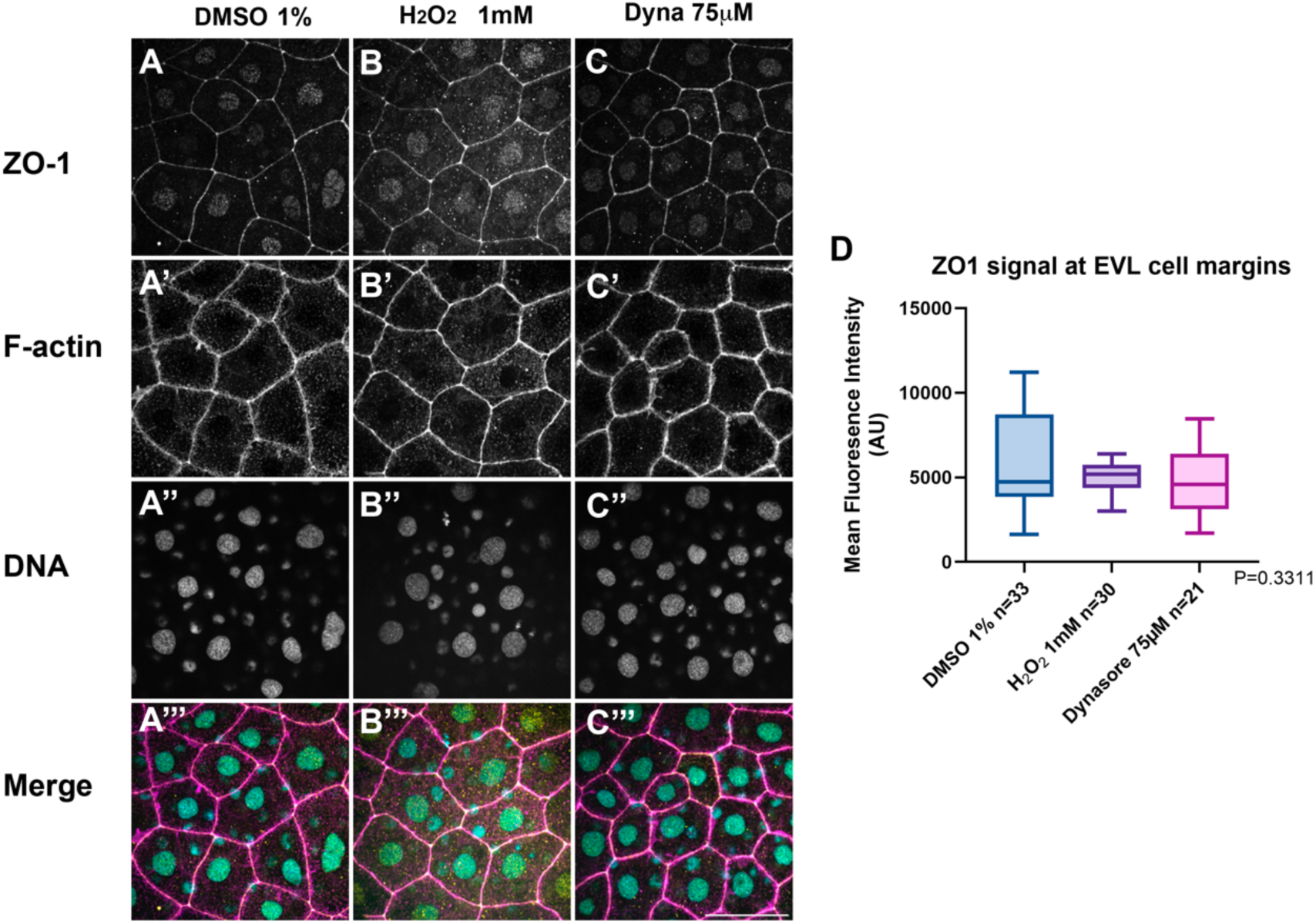
ZO-1 signal intensity at EVL cell margins is not affected by H_2_O_2_ and Dynasore treatment. All treatment groups started with embryos at oblong-sphere stage and the experiment stopped 3 hrs after the start of the treatment (around 60% epiboly). Embryos were then fixed, stained and visualized in a spinning disk confocal microscope. ZO-1 was visualized by immunolocalization in whole embryos in DMSO 1% treated EVL cells (A), H_2_O_2_ 1 mM (B), and 75 µM dynasore (C). Filamentous actin was visualized by Phalloidin Alexa Fluor 488 staining (A’-C’). Nuclei were visualized by staining DNA with DAPI (A’’-C’’). Images show merged ZO-1, F-actin and DNA channels (A’’’-C’’’). Scale bar, 50 µm. Quantification of ZO-1 signal at EVL cell margins (D). Values represent median and interquartile range; whiskers show minimum and maximum values. \**p* < 0.01 by nonparametric one-way analysis of variance (ANOVA) Kruskal Wallis with Dunns’s post hoc analysis (*n*, indicated in the graph). Enveloping layer (EVL). N=3 experimental repetitions.

**Supplementary Figure 7.**
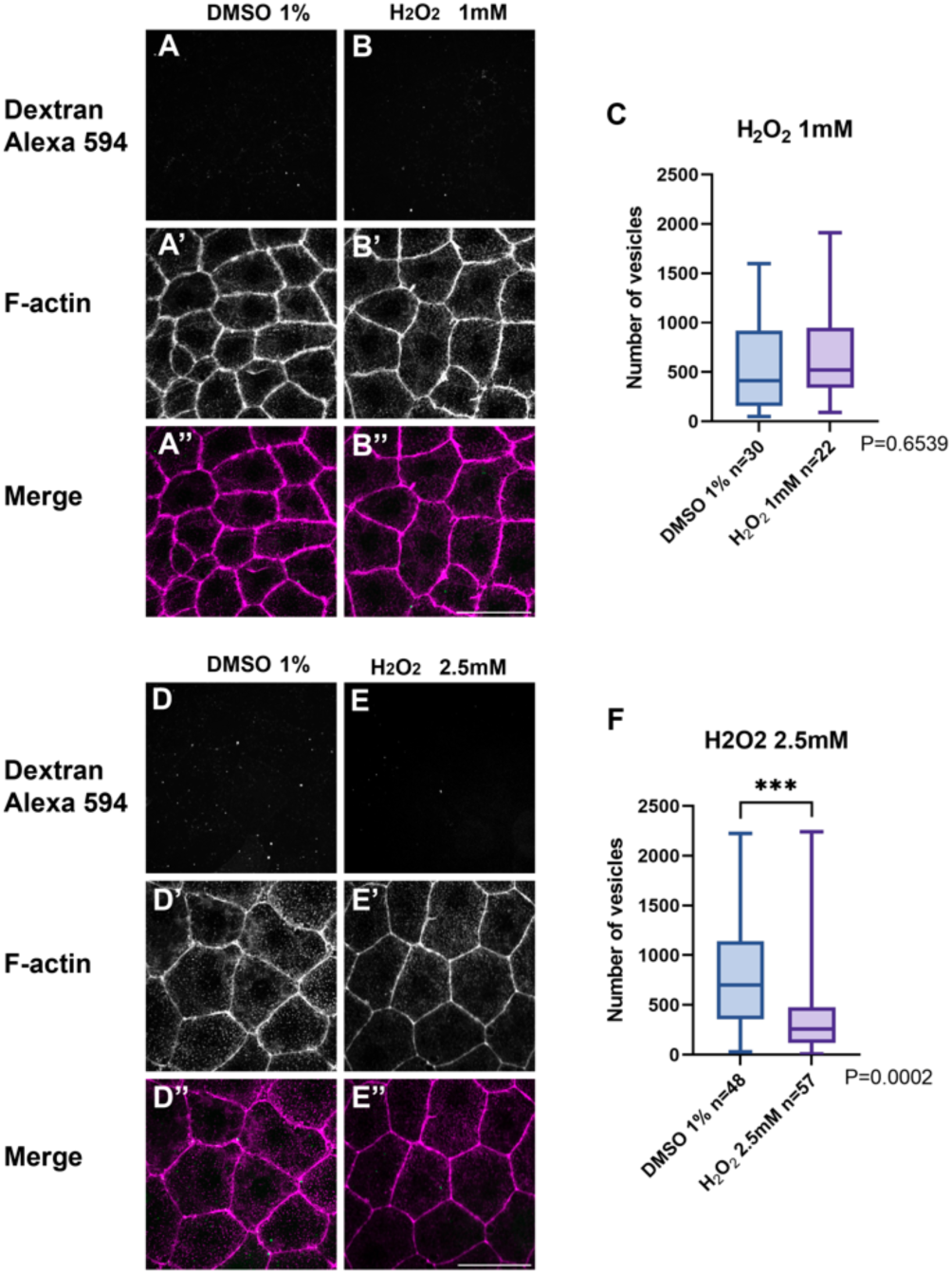
Increasing concentrations of H_2_O_2_ decreases fluid phase endocytosis at EVL cells. All treatment groups started with embryos at oblong-sphere stage and the experiment stopped 3 hrs after the start of the treatment (around 60% epiboly). Embryos were then fixed, stained and visualized in a spinning disk confocal microscope. Dextran fluorescent signal observed at EVL cells in control treated embryos (A), and H_2_O_2_ 1 mM (B) and H_2_O_2_ 2.5 mM (C). Filamentous actin was visualized by Phalloidin Alexa Fluor 488 staining (A’-C’). Scale bar, 50 µm. Quantification of the number of dextran vesicles at EVL cells (D and E). Values represent median and interquartile range; whiskers show minimum and maximum values. \**p* < 0.01 by nonparametric one-way analysis of variance (ANOVA) Kruskal Wallis with Dunns’s post hoc analysis (*n*, indicated in the graph).

**Supplementary Figure 8.**
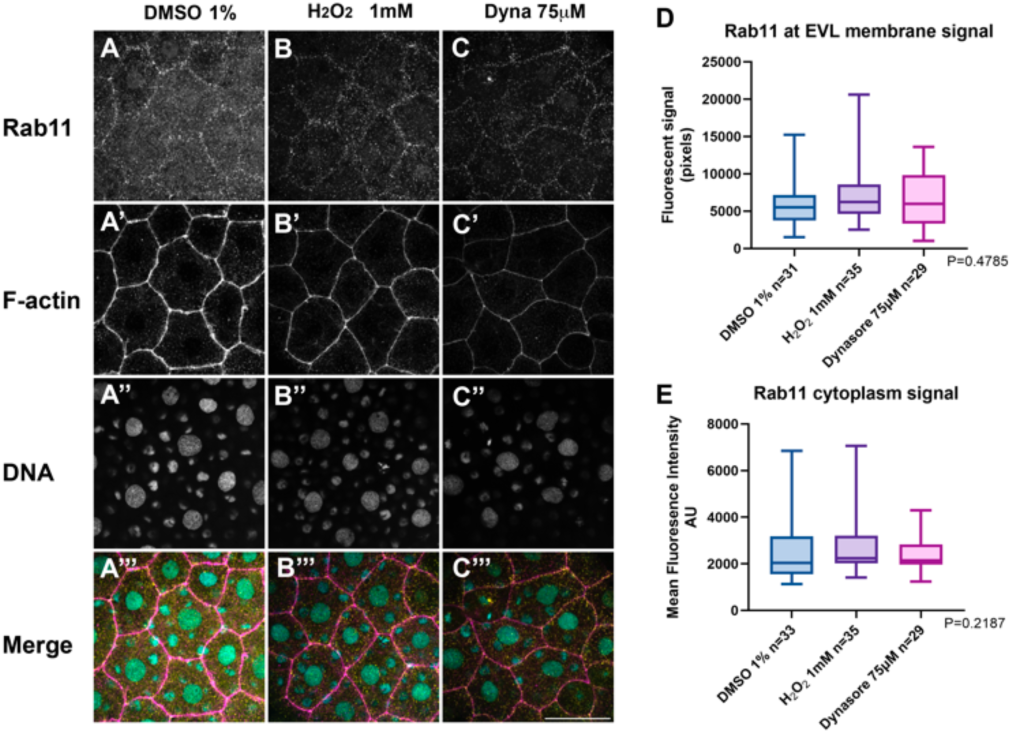
Rab11 signal intensity at EVL cell margins is not affected by H_2_O_2_ and Dynasore treatment. All treatment groups started with embryos at oblong-sphere stage and the experiment stopped when control treatment embryos reached around 65% epiboly. Embryos were then fixed, stained and visualized in a spinning disk confocal microscope. All images were obtained by spinning disk confocal microscopy. Rab11 was visualized by immunolocalization in whole embryos in DMSO 1% treated EVL cells (A), H_2_O_2_ 1 mM (B), and 75 µM dynasore (C). Filamentous actin was visualized by Phalloidin Alexa Fluor 488 staining (A’-C’). Nuclei were visualized by staining DNA with DAPI (A’’-C’’). Images show merged Rab11, F-actin and DNA channels (A’’’-C’’’). Scale bar, 50 µm. Quantification of Rab11 signal at EVL cell margins (D). *p* =0.4785. Quantification of Rab11 signal in the cytoplasm of EVL cells (E). *p* =0.2187. Values represent median and interquartile range; whiskers show minimum and maximum values. Statistical analysis with nonparametric one-way analysis of variance (ANOVA) Kruskal Wallis with Dunns’s post hoc analysis (*n*, indicated in the graph). N=3 experimental repetitions.

